# Rapid growth and defence evolution following multiple introductions

**DOI:** 10.1101/435271

**Authors:** Lotte A. van Boheemen, Sarah Bou-Assi, Akane Uesugi, Kathryn A. Hodgins

## Abstract

1. Rapid adaptation can aid invasive populations in their competitive success. Resource-allocation trade-off hypotheses predict higher resource availability or the lack of natural enemies in introduced ranges allow for increased growth and reproduction, thus contributing to invasive success. Evidence for such hypotheses are however equivocal and tests among multiple ranges over productivity gradients are required to provide a better understanding of the general applicability of these theories.
2. Using common gardens, we investigated the adaptive divergence of various constitutive and inducible defence-related traits between the native North American and introduced European and Australian ranges, whilst controlling for divergence due to latitudinal trait clines, individual resource budgets and population differentiation, using >11,000 SNPs.
3. Rapid, repeated clinal adaptation in defence-related traits was apparent despite distinct demographic histories. We also identified divergence among ranges in some defence-related traits, although differences in energy budgets among ranges may explain some, but not all, defence-related trait divergence. We do not identify a general reduction in defence in concert with an increase in growth among the multiple introduced ranges as predicted trade-off hypotheses.
4. *Synthesis*: The rapid spread of invasive species is affected by a multitude of factors, likely including adaptation to climate and escape from natural enemies. Unravelling the mechanisms underlying invasives’ success enhances understanding of eco-evolutionary theory and is essential to inform management strategies in the face of ongoing climate change.

## Introduction

Biological invasions are occurring at an accelerating pace due to the globalisation of anthropogenic activity (Ricciardi, 2007). Individuals colonizing new ranges likely face environments different from those previously experienced (Sax & Brown, 2000; Allendorf & Lundquist, 2003; Chown *et al.*, 2014). Nonetheless, alien populations often display enhanced performance compared to their native counterparts (Blossey & Notzold, 1995; Thébaud & Simberloff, 2001; Parker *et al.*, 2013), and this can be facilitated by rapid adaptation (Chown *et al.*, 2014; Colautti & Lau, 2015; Dlugosch *et al.*, 2015a). Therefore, in the face of ongoing environmental change, studies on introduced species are imperative to provide insight into invasive success as well as contemporary evolutionary processes.

Resource allocation trade-offs between life-history traits, such as growth rate and reproductive output, feature prominently in evolutionary theories developed to explain the success of invasive species (e.g., Hodgins & Rieseberg, 2011; Kumschick *et al.*, 2013; Turner *et al.*, 2014; Colautti & Lau, 2015). For instance, if abiotic stressors are mitigated upon introduction because of increased resource availability, increased investment in colonization or competitive ability could facilitate invasion success (Grime, 1977; Davis *et al.*, 2000; Bossdorf *et al.*, 2005; He *et al.*, 2010; Dlugosch *et al.*, 2015b). Similarly, the evolution of increased competitive ability hypothesis (EICA) postulates that release from specialist herbivores within the introduced range favours genotypes allocating resources to growth and reproduction in lieu of defence (Blossey & Notzold, 1995). Evidence for such adaptive divergence of invasive populations is however equivocal (Felker-Quinn *et al.*, 2013) perhaps due to allocation trade-offs among multiple competing functions (Mole, 1994; Züst & Agrawal, 2017), variation in resource availability or acquisition (Uesugi *et al.*, 2017; Züst & Agrawal, 2017), interplay with non-adaptive processes (Lee, 2002; Facon *et al.*, 2006; Prentis *et al.*, 2008; Rius & Darling, 2014; Estoup *et al.*, 2016), or other selective factors, such as climate, playing an important role in governing patterns of trait variation within and between ranges (Lachmuth *et al.*, 2011; Turner *et al.*, 2015).

Biotic and abiotic clines impacting plant resistance within ranges (Endara & Coley, 2011; Moles *et al.*, 2011a) can obscure the adaptive underpinnings of trait divergence governed by growth-defence trade-offs in response to changes during invasion in herbivory. For instance, herbivore pressure in the native range is expected to increase towards lower latitudes and potentially drive clines in plant defence in some species (Moles *et al.*, 2011a). This clinal pattern may be absent in the introduced range due to overall lack of herbivory, resulting in non-parallel defence gradients between ranges (e.g. Cronin *et al.*, 2015; Allen *et al.*, 2017). Moreover, high-resource environments support plant species with faster growth that are more vulnerable to herbivores (Coley *et al.*, 1985; Zandt, 2007; Endara & Coley, 2011), resulting in latitudinal clines in defence traits. Latitudinal clines in resource availability could subsequently lead to the evolution of high growth and reduced chemical defences at lower latitudes (Woods *et al.*, 2012; Moreira *et al.*, 2014), although this interspecific pattern may have limited application to intraspecific variation (Hahn & Maron, 2016, but see Woods et al., 2012). However, taken together these patterns suggest that the evolutionary consequences of herbivore escape could change along latitudinal gradients (Blumenthal, 2006). Geographical clines therefore need to be considered in tests of adaptive divergence between ranges (Colautti *et al.*, 2009).

The complex interplay between the evolutionary mechanisms shaping phenotypic divergence could also confound inferences of adaptation. Distinct demographic processes, including founder effects, genetic drift and admixture, often characterize introduction and alone can lead to divergence between native and introduced populations (Lee, 2002; Facon *et al.*, 2006; Prentis *et al.*, 2008; Rius & Darling, 2014; Estoup *et al.*, 2016). Dissection of the various evolutionary processes that can contribute to trait divergence is required to advance our understanding of rapid spread in invasive species. In addition, the repeatability of evolutionary patterns associated with introductions is unclear, as the majority of studies examining trait evolution following introduction focus on a single invaded range (e.g. Blossey & Notzold, 1995; Joshi & Vrieling, 2005; Hodgins & Rieseberg, 2011; Uesugi & Kessler, 2016, but see Colomer-Ventura *et al.*, 2015). Repeatable trait divergence across multiple invaded ranges would provide support for adaptive divergence of traits during in invasion as well as insight into selective mechanisms contributing to invasion success (Hodgins *et al.*, 2018; van Boheemen *et al.*, 2018).

The frequency and level of attack can impact the evolution of defence traits (Orrock *et al.*, 2015; Bixenmann *et al.*, 2016), which might also be expected to trade off due to their costs and redundancy (Koricheva *et al.*, 2004; Agrawal *et al.*, 2010). Predictable and strong attack should favour constitutive defence, whereas low, infrequent herbivory would favour no, or an inducible response (Agrawal & Karban, 1999; Ito & Sakai, 2009). These responses have been shown to vary over latitudinal clines within ranges (Moreira *et al.*, 2014 Rasmann & Agrawal, 2011). However, the studies exploring evolutionary shifts of constitutive and inducible defences between native and introduced ranges showed mixed results (e.g. Cipollini *et al.*, 2005; Eigenbrode *et al.*, 2008). Various Variable outcomes could result from a decrease in the intensity and frequency of herbivory following introduction (Maron & Vilà, 2001; Agrawal & Kotanen, 2003), including an increase in plasticity (Cipollini *et al.*, 2005; Lande, 2015) or high variability in inducible response among populations (Eigenbrode *et al.*, 2008). Testing such shifts in invasive species would provide insight into factors governing the evolution of induced/constitutive trait defence more generally.

*Ambrosia artemisiifolia* is a highly suitable system to study adaptive divergence in defence-related traits during invasion. This native North American weed has successfully established globally (Oswalt & Marshall, 2008), including recent introductions to Europe (~160 years ago Chauvel *et al.*, 2006) and Australia (~80 years ago; Palmer & McFadyen, 2012; van Boheemen *et al.*, 2017). Repeated clinal associations were found in *A. artemisiifolia* populations included in the current study, with declines in size and increase in SLA at higher latitudes (van Boheemen *et al.*, 2018), though differences occurred among ranges. At comparable latitudes, European plants were bigger and had lower SLA than natives, while Australian plants had higher SLA leaves (van Boheemen *et al.*, 2018).

We test quantitative trait divergence in 1) physical defence (trichome density), 2) chemical defence (phenolic compounds concentration and richness), and 3) inducibility of chemical defence among the native North American and introduced European and Australian ranges in a series of common garden experiments. Trichomes are found on the leaves and the stems of plants and deter herbivores (Kessler & Baldwin, 2002; Dalin *et al.*, 2008; Tian *et al.*, 2012). Phenolics are secondary metabolites that are often thought to confer resistance against herbivores (Bhattacharya *et al.*, 2010; War *et al.*, 2012; War *et al.*, 2018). These compounds are also known to be inducible in response to herbivore damage, as well as simulated herbivory treatments including wounding and methyl jasmonate (MeJA) applications (e.g. Lee *et al.*, 1997; Constabel & Ryan, 1998; Keinänen *et al.*, 2001; Heredia & Cisneros-Zevallos, 2009). We accounted for population structure, which could potentially drive patterns in traits that are non-adaptive, using >11,000 double-digest genotype-by-sequencing SNPs. Moreover, we controlled for defence-related trait variation along latitudinal clines.

We predict reduced constitutive defence within the introduced ranges together with elevated inducible response due to lower certainty of attack (Cipollini *et al.*, 2005) and a more plastic (inducible) response in recent colonisations (Lande, 2015). We expect non-parallel defence gradients between native and introduced ranges due to divergence of clines in herbivory (Moles *et al.*, 2011b) and/or variable resource gradients (Blumenthal, 2006; Hahn & Maron, 2016). Finally, we explored the association between defence-related trait divergence and divergence in growth and SLA among ranges as a growth-defence trade-off would result in greater growth in conjunction with reduced defence. However, greater defence could be facilitated by genotypes with enhanced resource acquisition resulting in a positive correlation in traits. By considering the complex interplay of the evolutionary mechanisms impacting defence divergence among multiple ranges, we test evolutionary changes in herbivore defence likely shaped by selection.

## Methods

### Study species

*Ambrosia artemisiifolia* is a highly invasive, monoecious, self-incompatible annual plant (Brandes & Nitzsche, 2006), most commonly found in disturbed habitats (Bassett & Crompton, 1975; Lommen *et al.*, 2017) and is expected to expand its range with ongoing climate change (Chapman *et al.*, 2014). It is the leading cause of hayfever worldwide (Taramarcaz *et al.*, 2005) and has a significant impact on crop yields (Kazinczi *et al.*, 2008). Within Europe, admixture following multiple introductions from distinct native sources has been suggested to have contributed to the success of introduced populations, and genetic variation equals levels observed in North America (Chun *et al.*, 2010; Gladieux *et al.*, 2010; Gaudeul *et al.*, 2011; van Boheemen *et al.*, 2017). A subsequent single bottlenecked introduction from Europe has been determined as the origin of the Australian invasion, although the exact European source is unknown (van Boheemen *et al.*, 2017).

Within the native range, around 450 herbivores have been associated with *Ambrosia* species, of which about 30% are specific to the *Ambrosia* genus (Gerber *et al.*, 2011). The North American native specialist *Ophraella communa* is shown to exert high levels of damage (Throop, 2005). Up to 50 polyphagous insect species have been associated with *A. artemisiifolia* in Europe, yet most cause little damage (Gerber *et al.*, 2011; Essl *et al.*, 2015). *Ophraella communa* has been sighted in Southern Switzerland and Northern Italy since 2013 (Müller-Schärer *et al.*, 2014), where it greatly affects *A. artemisiifolia* seedling survival and growth (Cardarelli *et al.*, 2018). In Australia, generalists *Zygogramma bicolorata* (leaf-feeding) and *Epiblema strenuana* (stem-boring) are widespread and seemingly exert some control (Palmer & McFadyen, 2012).

### Experimental set-up

To explore the divergence of constitutive quantitative defence traits between native and introduced ranges (“constitutive-defence experiment”), while accounting for divergence along latitudinal clines, we collected *Ambrosia artemisiifolia* seeds in 2013-2014 from broad geographical scales within the native North America and introduced Europe and Australia. We raised seedlings in a common garden (for detailed methods, see Supporting Information). Briefly, we stratified seeds for 6 weeks at 4°C (Willemsen, 1975). After a 2-week germination at 30°C with 12h light/dark cycle, we randomly transplanted the seedlings into 100ml kwikpot trays with Debco mix, followed by a second transplant to 0.7L pots containing Debco and 1.5ml slow-release fertilizer (Osmocote Pro, eight to nine months) one month later. We top-watered all plants and artificially manipulated daylight following the light cycle at the median latitude for all populations (47.3°N). To explore constitutive defence, we selected a seedling from four maternal lines, originating from 28 North American, 32 European and 20 Australian locations (Supporting Information, Table S1).

A separate greenhouse experiment was conducted to test whether the inducibility of defence response varied among plant origins (hereafter, “induction experiment”). We used a subset of populations used in the constitutive experiment (10 North American, 17 European and 12 Australian locations, Table S1). For each population, we selected four maternal lines, and grew two seedlings per line as above. One seedling per maternal line was allocated to either the control or simulated herbivory treatment. We simulated herbivory by vertically cutting off half of the newest fully formed leaf (wounding) and subsequently spraying the whole plant with 1mM methyl jasmonate (MeJA) (Campos-Vargas & Saltveit, 2002; Heredia & Cisneros-Zevallos, 2009; Hodgins & Rieseberg, 2011; Jordan *et al.*, 2015). Control plants were not wounded and were sprayed with distilled water.

### Trait measurements

For the constitutive experiment, we recorded trichome density at the mid-point of each plant under a dissecting microscope (Olympus, SZ-PT) using a 1 cm × 0.3 cm stem area at the mid-point of each plant, nine weeks after the second transplant. Three weeks later, we scanned one young, fully expanded leaf from each plant and calculated leaf area using ImageJ and the R package LeafArea (Katabuchi, 2015). We dried leaves at 45 °C for seven days and an addition 12 hours prior to weighing and weighed to the closest milligram. We calculated specific leaf area (SLA) by dividing leaf area by dry leaf weight (mm^2^/mg). We deconstructed plants for biomass measurements once the majority of seeds had ripened. We placed aboveground components in paper bags and dried these in ovens at 45 °C for at least 36 hours. Before dry weight biomass measures, we dried materials for an additional minimum of 24 hours to ensure the dry weight was constant at the time of measuring and it was not variable due to humidity in the air or incomplete drying. We weighed this shoot biomass to the closest 0.1 gram.

Leaf samples for phenolic analyses were collected four weeks after the second transplant by clipping approximately 200 mg of the newest fully expanded leaf, which was flash frozen in liquid nitrogen and stored in a −80 °C. In the induction experiment, we collected leaf samples 24 hours after the final treatment. Samples were extracted in 1 ml of 80% methanol (% by volume in water) using a Qiagen TissueLyser II for 30 seconds at 30 rps twice and centrifuged for 30 minutes at 2700 rpm. Phenolic samples from the constitutive-defence experiment were analysed using HPLC Agilent 1200 series (Agilent Technologies Australia, Mulgrave, VIC, Australia) equipped with C18 reverse-phase column (Waters, 5.0 μm, 250 mm × 4.6 mm; Alltech Australia, Baulkham Hills NSW, Australia). The elution system consisting of solvents (A) 0.25% H3PO4 in water (pH 2.2) and (B) acetonitrile was: 0–6 min, 0–12% of B; 6–10 min, 12-18% of B, and 10-30 min, 18-58% of B, with a flow rate of 1 mL/min and injection volume of 15 µL (Keinänen *et al.*, 2001). Samples from the induction experiment were analysed with Agilent Infinity 1260 equipped with C18 reverse-phase column (Poroshell 120 EC-C18, 2.7 μm, 150 mm × 3.0 mm; Agilent Technologies Australia, Mulgrave, VIC, Australia). The elution method was modified from above and was: 0–2 min, 0–12% of B; 2–3.3 min, 12–18% of B, and 3.3–10 min, 18-58% of B, with a flow rate of 0.5 mL/min and injection volume of 5 µL. In both experiments, phenolic compound peaks were identified to their compound classes using UV spectra and relative abundance was quantified at 320 nm. To estimate phenolic compound richness, we counted the number of detectable peaks. The relative concentration of eight major phenolic peaks was estimated as area under each peak divided by sample fresh weight. Results could not be directly compared as the two experiments were performed in different greenhouses and samples from each experiment were run using different HPLC machines.

### Statistical analyses

To test if constitutive defence differed among ranges (the constitutive experiment), we examined individual phenolic compound composition in a multivariate analysis of covariance (MANCOVA) and the concentration of individual phenolic compounds, phenolic compound richness, total phenolic concentration, and trichome density in univariate mixed models. To account for latitudinal variation within and among ranges, each multi- and univariate model included range, latitude, their interaction and a latitude^2^ effect as fixed factors. To control for neutral population structure, possibly shaping trait variation between populations, univariate models included q-values as a random effect, as obtained from STRUCTURE analysis performed on genetic data. For the multi- and univariate analyses, we improved normality of the data by square-root- or log-transforming traits where appropriate. For the MANCOVA, we included the concentration of eight major phenolic compounds (Spearman’s ρ among peaks < 0.75) and calculated Wilks’ λ (multivariate F-value) to measure the strength of the associations. To measure the variance explained by the fixed effects or the full model within the univariate models, we calculated the marginal and conditional coefficients of determination using the MuMIn package (Bartón, 2018). We computed type III Wald F-values with Kenward-Roger degrees of freedom and step-wise removed non-significant effects, starting with the highest order interaction. For univariate models, we plotted the partial residuals of each response variable by ranges, thus accounting for latitudinal clines and neutral population genetic structure and reported these adjusted means and standard errors for each range, calculated using the *phia* packaged (De Rosario-Martinez, 2015).

To explore the variation in inducibility among ranges (the induction experiment), we repeated the steps for the constitutive experiment, now including treatment and its interactions with range and latitude as fixed effects. For the MANCOVA, we included five peaks (excluding three with Spearman’s ρ > 0.75) to increase power of this test (Scheiner, 2001). We retained treatment in these models, as this was the variable of interest. Here, a significant treatment effect would signify an inducible response, whereas a treatment × range interaction would imply this response differs between ranges. A treatment × latitude interaction would indicate different inducibility at different latitudes. To test if variation in induction differed between ranges (Eigenbrode *et al.*, 2008), we compared the coefficient of variation (c_v_) using the modified signed-likelihood ratio test for equality with 10^4^ simulations in the cvequality package (Krishnamoorthy & Lee, 2014; Marwick & Krishnamoorth, 2018).

To examine associations between defence-related traits and plant growth and to assess if divergence in individual resource budgets could have resulted in range differences in defence-related trait investment, we tested responses of phenolic richness, phenolic concentrations or trichome density to shoot biomass or SLA. Each model included a defence-related trait as response, with shoot biomass or SLA, range and their interaction as predictors. We used individual trait values and included individual STRUCTURE q-values and sampling location as random factors. We explored significant range × defence interactions using a Holm p-value correction in the *phia* package (De Rosario-Martinez, 2015). In these models, a negative association between defence-related traits and shoot biomass would suggest a trade-off, while a positive one might indicate differences in resource acquisition. Range differences at similar values of shoot biomass or SLA would indicate defence-related trait divergence independent of genotypic differences in individual resource budgets.

To explore if constitutive and inducible defence trade off, we first calculated the induced level of total phenolics for each maternal line as the difference between damage and control treatments of the two half-sibs. This estimate of induction is thought to reduce correlations with control treatment estimates and thus the collinear associations (e.g. due to genotypic biases) will not mask the trade-off associations (Morris *et al.*, 2006). We included population of origin and individual q-values as random factor in these models. A significant negative association between induced and constitutive levels of phenolic concentration and richness would indicate the presence of a trade-off. All statistical analyses were conducted in R v3.4.3 (R Core Team, 2018).

## Results

### Constitutive defence trait divergence between ranges

We found significant range divergence in constitutive phenolic composition (Table 1a), resulting from differences between the introduced Europe and the native North America (F_8,66_=3.280, p=0.010, Wilks’ λ=0.716) (Table 1b). Total phenolic concentration was similar between the native and European populations, but 28% lower in Australia (Table 1b&c, Fig. 1). Phenolic peak richness differed among ranges (Table 1): it was highest in the introduced European range (adjusted mean of 43 peaks) followed by the native North American (40 peaks) and introduced Australian ranges (33 peaks). Trichome density showed no differences between ranges (Table 1a, Fig. 1). The composition of individual phenolic compounds and peak richness depended on latitude, though no such effect was found for the total phenolic concentration or trichome density (Table 1a, Fig. 2). We did not observe range × latitude interactions for any of the defence-related traits (Table 1a), suggesting latitudinal clines, when present, did not differ between ranges.

**Table 1.**
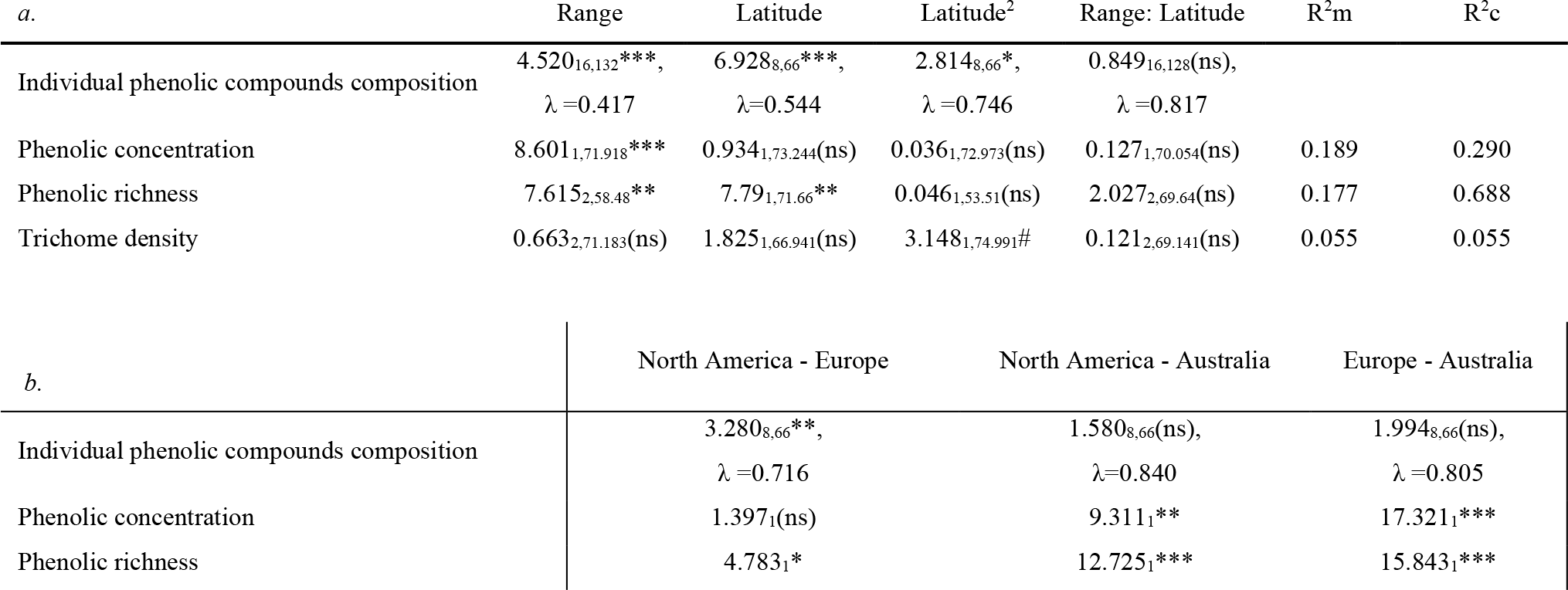
*Ambrosia artemisiifolia* defence-related trait responses (population means) to range, latitude, their interaction and latitude^2^ (to account for non-linear relationship) in the constitutive experiment in multivariate (individual phenolic compounds) and univariate analyses (a), with dissection of significant range effects (p<0.05) in post-hoc tests (b). We reported Wald type III F (a) or χ^2^ test values (b), Kenward-Roger degrees of freedom (subscript), significance (symbols). In the multivariate analysis, Wilk’s λ measure the strength of the association, in univariate analyses, marginal (R^2^m) and conditional (R^2^c) coefficients measure the variance explained by fixed effects or full models. Models were step-wise reduced starting with the highest order non-significant interaction and univariate analyses included neutral population genetic structure as a random effect. *ns: p*>0.1; #*: p*<*0.1*, \**: p*<*0.05*; \*\**: p*<*0.01*; \*\*\**: p*< *0.001*

**Fig. 1.**
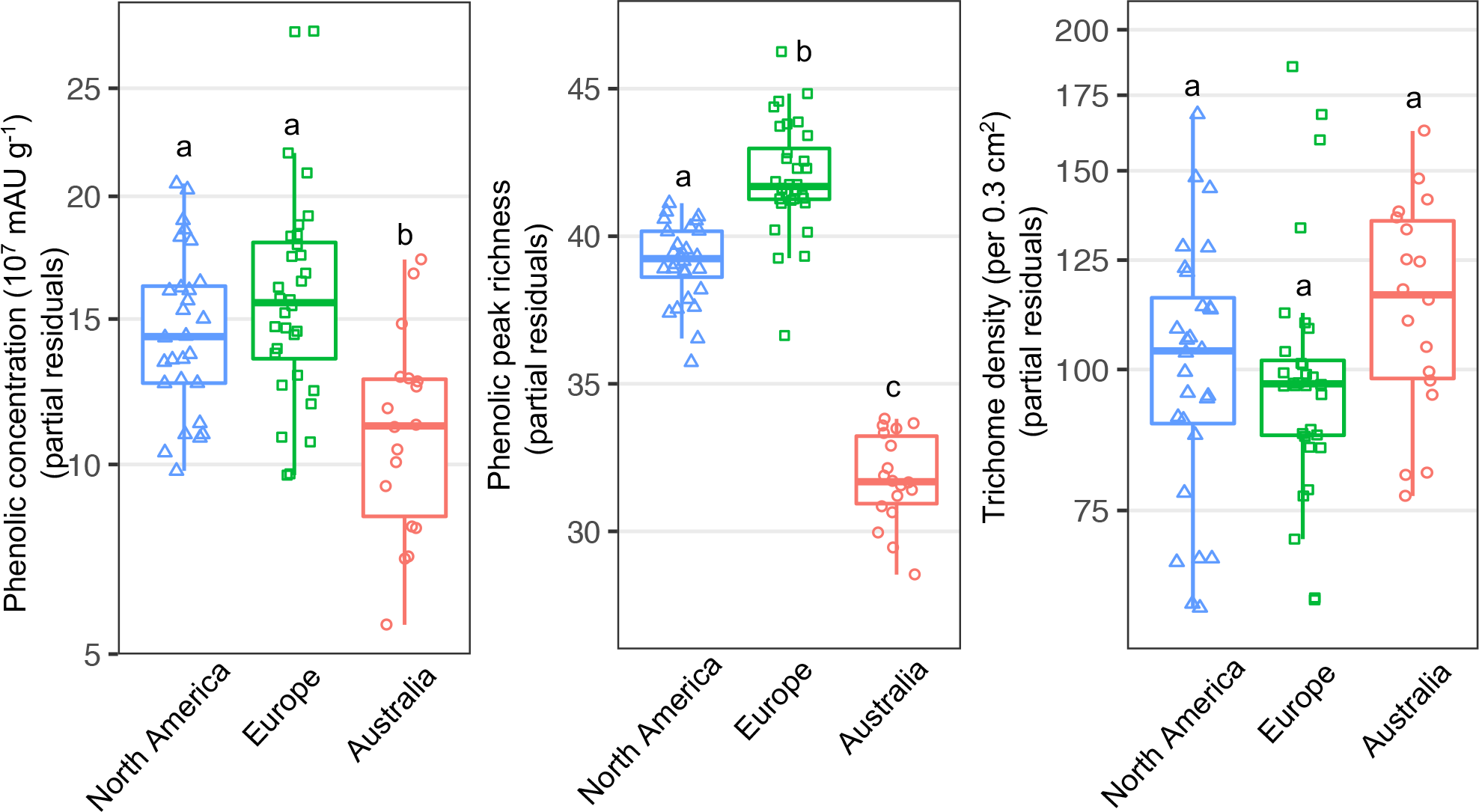
Partial residual defence trait responses (phenolic concentration, peak richness and trichome density) of *A. artemisiifolia* populations to range, accounting for latitudinal clines and neutral population structure. Different letters indicate significance for pairwise range comparisons (Table 1).

**Fig. 2.**
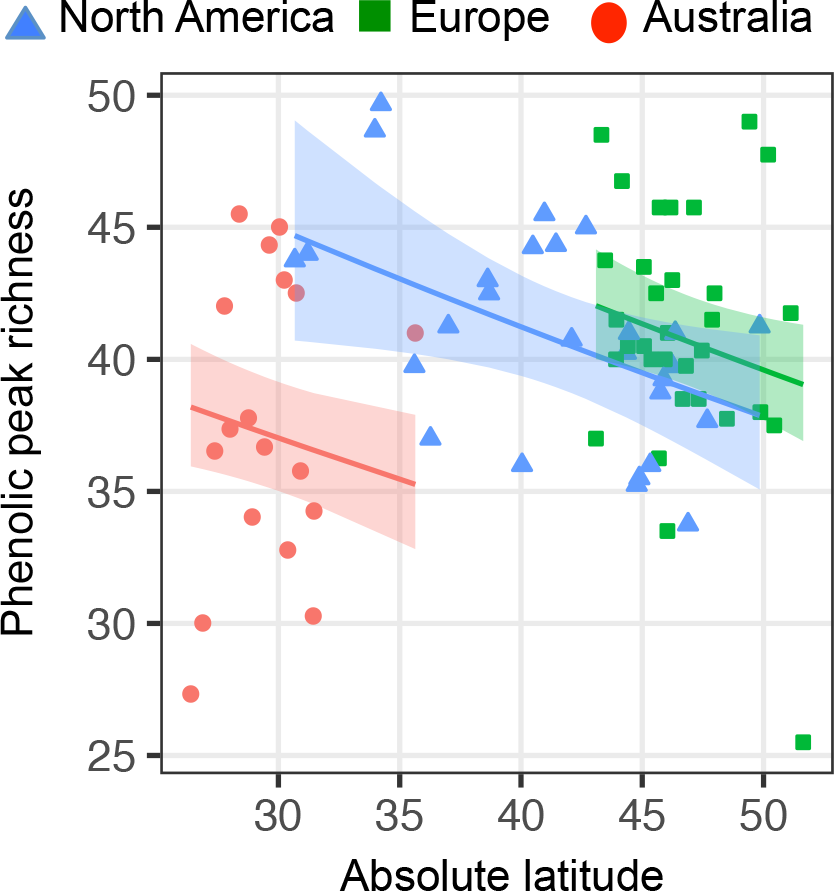
Population mean response of phenolic peak richness to range (native North America, blue triangles; introduced Europe, green squares; introduced Australia, red circles) and latitude in *Ambrosia artemisiifolia*, with predicted latitudinal clines (+/− 95% confidence interval) corrected for neutral population structure.

### Inducible defence trait divergence between ranges

We found a significant treatment effect on individual phenolic compound composition in the induction experiment (F_5,59_=12.014, p<0.001, Wilks’ λ=0.496) (Table 2). The total phenolic concentration was slightly suppressed in the herbivory-simulating treatment (Table 2, Fig. 3), but the phenolic peak richness did not show a response to experimental treatment (Table 2, Fig. 3). We identified no treatment × range × latitude interactions (Table 2, Fig. 3), suggesting there is no range difference in inducibility clines. Also, the absence of treatment × latitude interactions (Table 2, Fig. 3), suggests an overall lack of latitudinal clines in inducibility. Moreover, no treatment × range interactions (Table 2, Fig. 3) suggests the inducible response did not differ between ranges. We did not find range differences in the variation of inducible phenolic peak richness (c_v_=1.401, p=0.496) or concentration (c_v_=2.297, p=0.317).

**Table 2.** *Ambrosia artemisiifolia* defence-related trait responses (population means) to range, latitude, treatment, their interactions and latitude^2^ in the inducible experiment in multivariate (individual phenolic compounds) and univariate analyses. Range, latitude, their interaction or latitude^2^ were included as covariates and significant results were not explored further. We reported Wald type III F, Kenward-Roger degrees of freedom (subscript), significance (symbols) (a). In the multivariate analysis, Wilk’s λ measure the strength of the association, in univariate analyses, marginal (R^2^m) and conditional (R^2^c) coefficients measure the variance explained by fixed effects or full models (a). Models were step-wise reduced starting with the highest order non-significant interaction and univariate analyses included neutral population genetic structure as a random effect. *ns: p>0.1; *: p<0.05; **: p<0.01; ***: p< 0.001*

|  | Range | Latitude | Latitude<sup>2</sup> | Range:Latitude | Treatment | Treatment:Range | Treatment:Latitude | Treatment:<br>Range:<br>Latitude | R<sup>2</sup><sub>m</sub> | R<sup>2</sup><sub>c</sub> |
| --- | --- | --- | --- | --- | --- | --- | --- | --- | --- | --- |
| Individual phenolic<br>compounds concentration | 7.591<sub>10,118</sub>***,<br>$\lambda=0.370$ | 10.637<sub>5,59</sub>***,<br>$\lambda=0.526$ | 4.818<sub>5,59</sub>**,<br>$\lambda=0.710$ | 2.311<sub>10,118</sub>*,<br>$\lambda=0.699$ | 12.014<sub>5,59</sub>***,<br>$\lambda=0.496$ | 0.326<sub>10,112</sub>(ns),<br>$\lambda=0.944$ | 0.977<sub>5,58</sub>(ns),<br>$\lambda=0.922$ | 1.357<sub>10,108</sub>(ns),<br>$\lambda=0.789$ | | |
| Phenolic concentration | 4.970<sub>2,30.905</sub>* | 5.932<sub>1,31.505</sub>* | 1.577<sub>2,28.556</sub>(ns) | 1.745<sub>1,31.505</sub>(ns) | 4.241<sub>1,35.628</sub>* | 0.005<sub>2,32.285</sub>(ns) | 1.077<sub>1,35.428</sub>(ns) | 1.417<sub>2,31.192</sub>(ns) | 0.201 | 0.457 |
| Phenolic richness | 3.764<sub>2,29.93</sub>* | 4.700<sub>1,31.08</sub>* | 1.340<sub>2,28.24</sub>(ns) | 6.030<sub>1,30.95</sub>* | 0.850<sub>1,35.33</sub>(ns) | 0.091<sub>2,32.1</sub>(ns) | 0.825<sub>1,34.89</sub>(ns) | 1.923<sub>2,30.78</sub> (ns) | 0.323 | 0.723 |

**Fig. 3.**
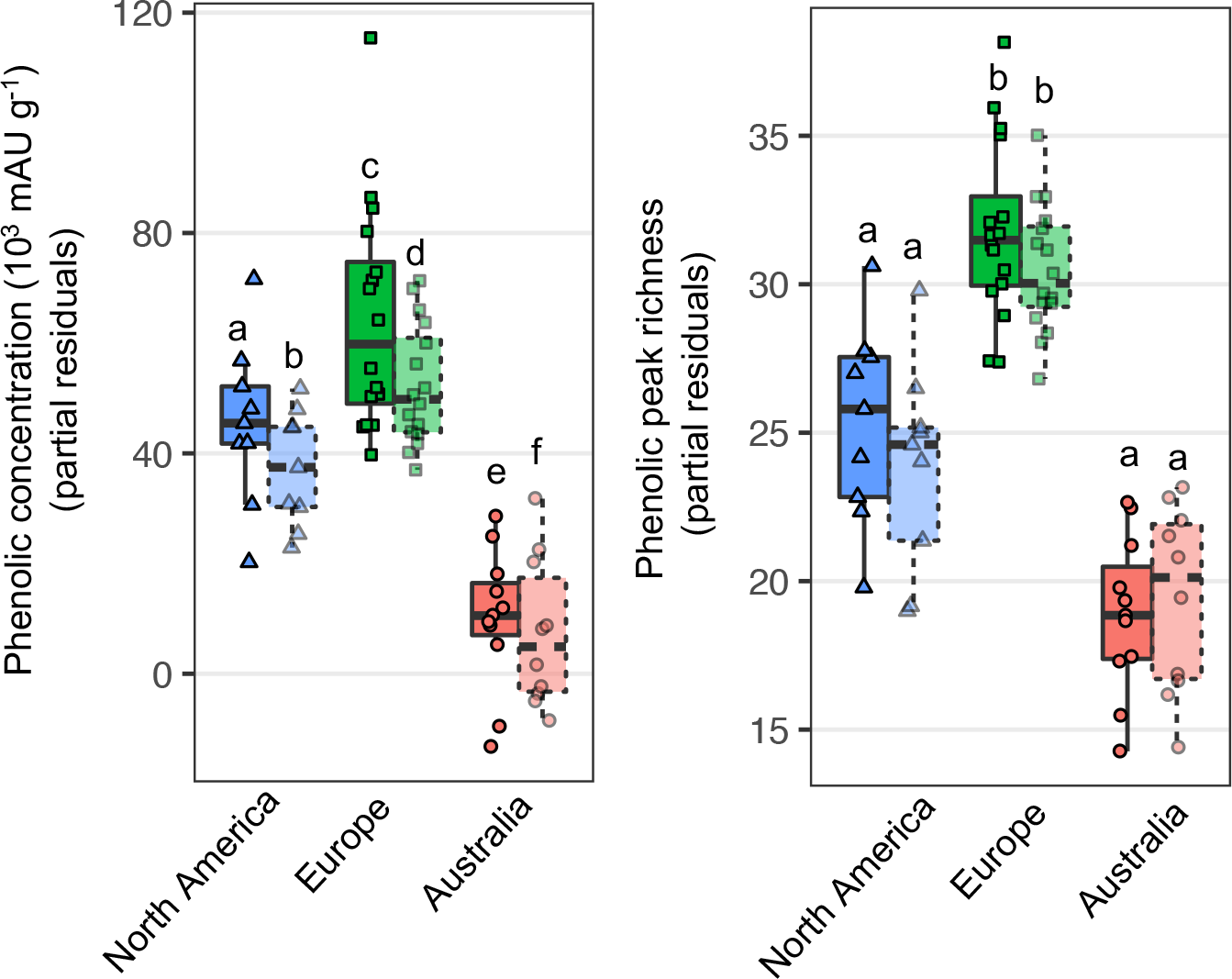
Partial residual defence trait responses (phenolic concentration and peak richness) of *A. artemisiifolia* populations to control (solid symbols) and herbivore simulating treatment (wounding + MeJA, dashed transparent symbols), with covariates of range, accounting for latitudinal clines and neutral population structure. Letters indicate significance of effect (Table 2).

### Associations between defence, biomass and specific leaf area (SLA)

Within each range, phenolic concentration and richness was positively correlated with shoot biomass, whereas we found a negative association between trichome density and shoot biomass (Table 3, Fig. 4). We found high-SLA leaves had lower phenolic concentration and peak richness, yet higher trichome density (Table 3, Fig. 4). No interactions were significant between range and predictor variables (shoot biomass or SLA), suggesting these associations among traits were consistent between ranges (Table 3, Fig. 4). These results emphasize the close relationship between plant growth, physiology and defence.

**Table 3.** Constitutive defence trait response of *Ambrosia artemisiifolia* individuals to shoot biomass, specific leaf area and their interaction with range (a), with dissection of significant range effects (p<0.05) in post-hoc tests (b). We reported corresponding figure, Wald type III F (a) or χ^2^ test values (b), Kenward-Roger degrees of freedom (subscript) and significance (symbols). Marginal (R^2^m) and conditional (R^2^c) coefficients measure the variance explained by fixed effects or full models (a). Models were step-wise reduced starting with the highest order non-significant interaction and included population origin and neutral population genetic structure as random effects. *ns: p*>*0.1*; #*: p*<*0.1*, \**: p*<*0.05*; \*\**: p*<*0.01*; \*\*\**: p*< *0.001*

| Predictor | Response | Figure 4 | Range | Predictor | Range:Predictor | R<sup>2</sup>m | R<sup>2</sup>c |
| --- | --- | --- | --- | --- | --- | --- | --- |
| Shoot biomass | Phenolic concentration | A | 19.321<sub>2,79.435</sub>*** | 31.098<sub>1,181.299</sub>*** | 1.441<sub>2,184.26</sub>(ns) | 0.160 | 0.189 |
|  | Phenolic richness | B | 18.96<sub>2,79.29</sub>*** | 53.389<sub>1,180.51</sub>*** | 1.565<sub>2,187.82</sub>(ns) | 0.199 | 0.224 |
|  | Trichome density | C | 10.242<sub>2,79.4</sub>*** | 18.49<sub>1,174.06</sub>*** | 0.525<sub>2,180.84</sub>(ns) | 0.093 | 0.103 |
| Specific leaf area | Phenolic concentration | D | 6.162<sub>2,71.167</sub>** | 38.464<sub>1,208.912</sub>*** | 0.98<sub>2,202.53</sub>(ns) | 0.202 | 0.269 |
|  | Phenolic richness | E | 5.349<sub>2,69.16</sub>** | 42.692<sub>1,217.97</sub>*** | 0.32<sub>2,202.6</sub>(ns) | 0.215 | 0.331 |
|  | Trichome density | F | 1.828<sub>2,71.7</sub>(ns) | 10.994<sub>1,204.66</sub>** | 1.196<sub>2,206.12</sub>(ns) | 0.049 | 0.121 |

b.
| Predictor | Response | Figure 4 | North America - Europe | North America - Australia | Europe - Australia |
| --- | --- | --- | --- | --- | --- |
| Shoot biomass | Phenolic concentration | A | 5.846<sub>1</sub>* | 22.155<sub>1</sub>*** | 40.087<sub>1</sub>*** |
|  | Phenolic richness | B | 2.546<sub>1</sub>(ns) | 26.667<sub>1</sub>*** | 37.964<sub>1</sub>*** |
|  | Trichome density | C | 2.255<sub>1</sub>(ns) | 13.049<sub>1</sub>*** | 21.047<sub>1</sub>*** |
| Specific leaf area | Phenolic concentration | D | 2.152<sub>1</sub>(ns) | 5.032<sub>1</sub>* | 12.485<sub>1</sub>** |
|  | Phenolic richness | E | 0.459<sub>1</sub>(ns) | 6.655<sub>1</sub>* | 10.643<sub>1</sub>** |
|  | Trichome density | F | - | - | - |

**Fig 4.**
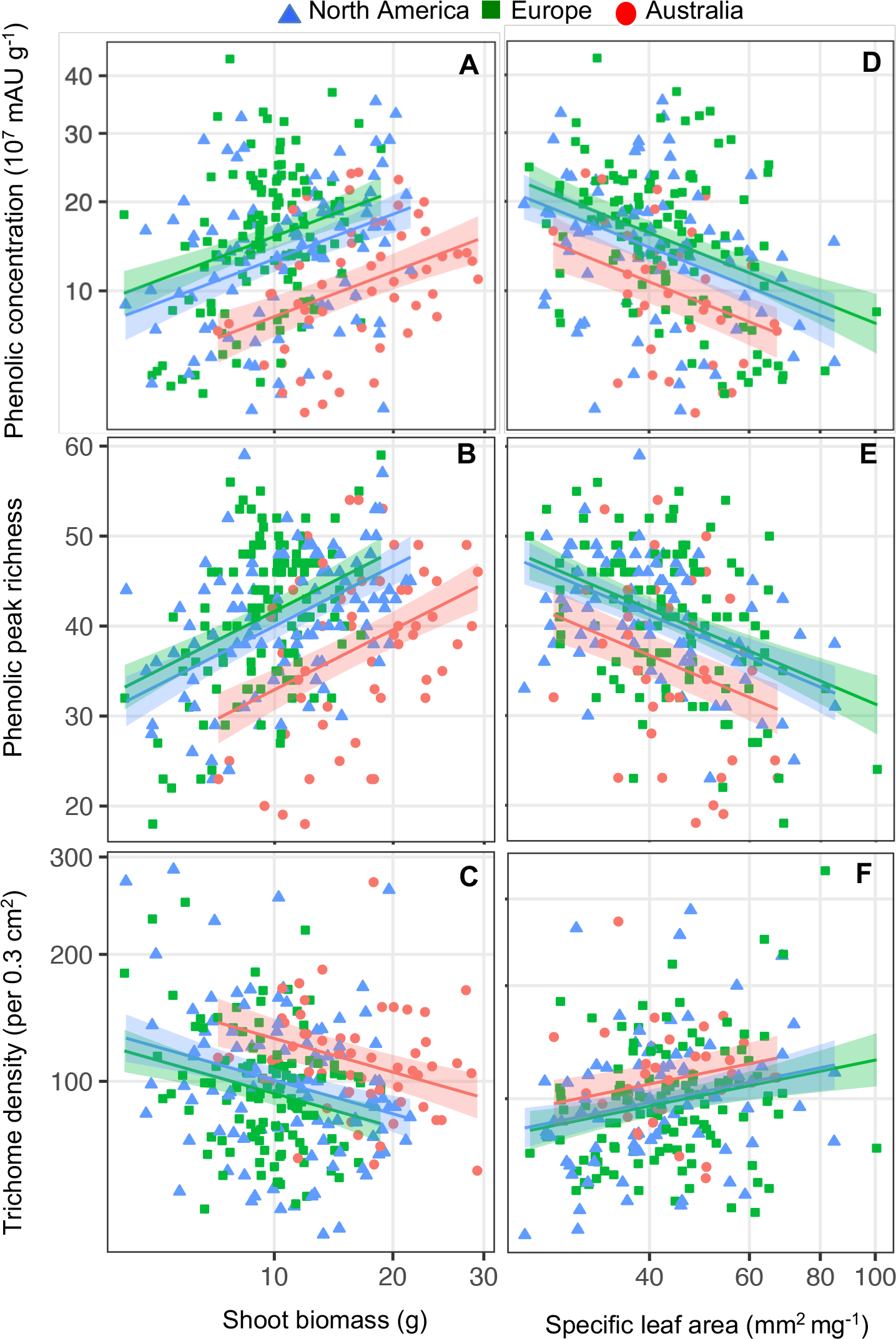
Defence trait responses (phenolic concentration, peak richness and trichome density) of *A. artemisiifolia* individuals to range (native North America (blue triangles); Europe (green squares); Australia (red circles)), shoot biomass (left panels) or specific leaf area (right panels) with model predictions (+/− 95% confidence interval, Table 3).

When controlling for shoot biomass or SLA, total phenolic concentration in European plants was higher compared to North American individuals of comparable weight (Table 3, Fig. 4), whereas no difference existed in latitude models (Table 1, Fig. 1). Conversely, phenolic peak richness was no longer significantly different between North America and Europe (Table 3, Fig. 4) compared to range comparisons accounting for latitude (Table 1, Fig. 1). Australian plants exhibited lower phenolic concentration and peak richness compared to native or European plants of comparable weight or SLA. Yet, at the same plant weight, Australian plants had higher trichome densities than in the other ranges (Table 3, Fig. 4). These patterns match previous analyses including latitude (Table 1, Fig. 1).

### Constitutive-inducible trade-offs

Induced levels of phenolic concentration and richness, the response variables, were negatively associated with the predictors, the constitutive levels (concentration: F_1,141.88_=76.286, p<0.001; richness: F_1,123.81_=78.126, p<0.001; Fig. 5). We found no range differences in either induced response trait (concentration: F_2,31.07_=0.265, p=0.769; richness: F_2,31.518_=1.719, p=0.196; Fig. 5), nor did we identify interactions between range and the predictor constitutive phenolic concentration (F_2,141.11_=0.866, p=0.423). Range × constitutive phenolic peak richness (F_2,129.82_=3.045, p=0.051) was marginally significant. These results suggest that constitutive and inducible defence trade off, although there is no difference between ranges.

**Fig. 5.**
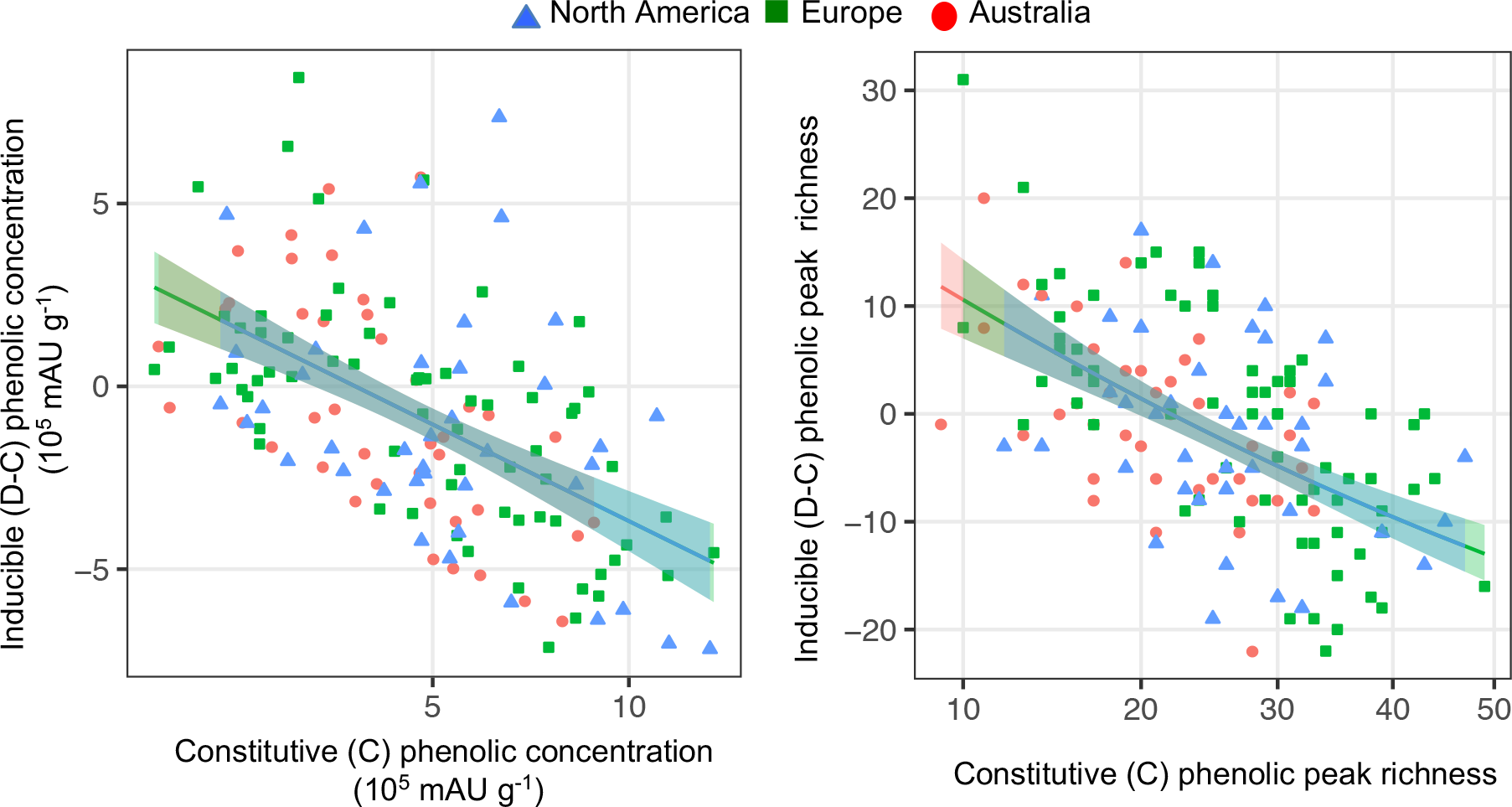
Inducible (D: wounding + MeJA; C: control) versus constitutive (control) defence trait responses (phenolic concentration and peak richness) of *A. artemisiifolia* populations among ranges (native North America: blue triangles; Europe: green squares; Australia: red circles) with model predictions (+/− 95% confidence interval).

## Discussion

In this study, we found evidence for divergence in defence related traits within and between ranges. Repeated latitudinal clines in phenolic richness and individual phenolic composition were identified, suggesting rapid adaptation of phenolics to local environments following invasion. Though we observed reduced phenolic concentration and richness in introduced Australia compared to the native plants while controlling for genetic structure, levels were similar or slightly higher in the introduced Europe compared to native populations at comparable latitudes and energy budgets. In addition, trichome density did not differ among ranges. These patterns are inconsistent with the Evolution of Increased Competitive Ability (EICA) hypothesis. In line with predictions, however, a trade-off between the constitutive and inducible phenolics was observed together with similar phenolic inducibility among ranges. To our knowledge, this is the first study testing the evolution of defence related traits across multiple introductions while exploring the predicted confounding of latitudinal clines, population substructure or genotypic differences in individual resource budgets. Therefore, the apparent absence of the predicted repeated selection against high defence investment following introduction is unlikely to be entirely masked by these factors. We examine these processes in detail and suggest alternative mechanisms driving defence-trait divergence within and among native and introduced ranges.

### Divergence in constitutive defence-related traits

The rapid and repeated latitudinal divergence in phenolic compound composition and richness populations suggests direct or indirect selection of latitude-associated factors. Corresponding to our findings, typical reported patterns include high growth and low defence at more productive high-resource (Coley *et al.*, 1985; Zandt, 2007; Endara & Coley, 2011), low-latitude environments (Woods *et al.*, 2012; Moreira *et al.*, 2014; Hahn & Maron, 2016, Blumenthal, 2006). Native clines in herbivore load could result in such observations, though the predicted herbivore reduction following introduction should lead to non-parallel defence clines among native and introduced ranges (Cronin *et al.*, 2015; Allen *et al.*, 2017). However, in our data, latitudinal clines in defence-related traits (phenolic compound composition and peak richness) were parallel, which could reflect consistent patterns of selection with latitude in all three ranges. The absence of the predicted patterns could result from parallel clines in herbivore loads in each range or the presence of alternative evolutionary forces driving latitudinal trait divergence in the multiple ranges. Indeed, clinal variation in herbivory is not as common as previously thought (Moles *et al.*, 2011a)), although geographic information on *A. artemisiifolia* herbivore pressure is needed.

Alternatively, latitudinal clines could arise through direct selection on the alternative functions of phenolic compounds, or indirect selection through genetic covariance with traits under climate mediated selection. Climate was previously shown to be a more important driver of trait divergence compared to enemy release (Colautti *et al.*, 2009; Colautti & Barrett, 2013; Colomer-Ventura *et al.*, 2015). Along these lines, climatic differences between the ranges not captured by latitude could contribute to patterns of divergence in Australian defence-related traits. For instance, trichomes protect plants from UV (Bassman, 2004; Hauser, 2014) and selection for this alternate function in high-UV Australia (WHO, 1998) could potentially explain the higher density of trichomes in this range when controlling for plant size. Herbivore exclusion experiments at various latitudes and environments would be important for to disentangling how resource availability, herbivory and other climatic factors might interact during invasion and impact the evolution of growth and defence traits.

When correcting for these latitudinal clines, we found conflicting patterns of defence-related trait divergence between the native and two introduced ranges. Genotypic differences in resource acquisition (Van Noordwijk & de Jong, 1986; Agrawal, 2011; Züst & Agrawal, 2017) and historical contingency (Lee, 2002; Facon *et al.*, 2006; Prentis *et al.*, 2008; Rius & Darling, 2014; Estoup *et al.*, 2016) can obscure trade-offs predicted under resource-allocation trade-off hypotheses. Accordingly, we show trichome density, phenolic concentration and peak richness were strongly associated with plant biomass and specific leaf area (SLA)(Fig. 4). Contrary to EICA predictions, phenolic peak concentration was significantly higher in Europe compared to native North America at comparable shoot biomass, although this difference disappeared when controlling for latitude or SLA. Similarly, phenolic richness was significantly higher in Europe than North America at equivalent latitudes, but this likely reflects the larger size and lower SLA of European plants at similar latitudes (van Boheemen *et al.*, 2018). However, lower phenolic concentration and peak richness in Australia was still present at similar latitude, biomass or SLA compared to North America. Invasion history is unlikely a major factor in this observed defence-related trait divergence as we accounted for population genetic structure in our analysis.

An adaptive reduction of constitutive defence traits following introduction to Europe and Australia was predicted due to a general release from natural enemies. However, levels of chemical defence-related traits (phenolic concentration and richness) were not consistently lower in introduced ranges compared to native populations. Such unexpected findings could have resulted from variation in contemporary herbivory among introduced ranges. Of particular relevance to the EICA hypothesis are specialist herbivores, as herbivory by specialists, but not necessarily generalists, is hypothesized to consistently decline during invasion (Müller-Schärer *et al.*, 2004; Joshi & Vrieling, 2005; Felker-Quinn *et al.*, 2013). Indeed, introduced Japanese *A. artemisiifolia* populations re-exposed to specialist leaf beetle *Ophraella communa* for >10 years were more resistant than herbivore-free populations (Fukano & Yahara, 2012). However, rapid adaptation to *O. communa* is unlikely to have led to the observed elevated European phenolic concentration and richness, as the seeds used in our experiment were collected in 2014 and this beetle is constrained to southern Europe since introduction in 2013 (Sun *et al.*, 2017).

Alternatively, differences in generalist load between introduced ranges could have resulted in variation in quantitative digestibility-reducing chemicals (e.g., phenolics), which defend against both generalist and specialists (Müller-Schärer *et al.*, 2004). Surveys describe a high diversity of generalist species in Europe (Gerber *et al.*, 2011; Essl *et al.*, 2015) but not in Australia (Palmer & McFadyen, 2012) suggesting herbivory in this species is higher in Europe than Australia. However, Genton *et al.* (2005) previously found that compared to native Ontario, the most common forms of damage (chewing and perforation) together with the generalist herbivore load was reduced in introduced France populations consistent with enemy escape in Europe compared to native North America. Contradicting EICA expectations, but consistent with our findings for Europe, the French plants showed no evolutionary loss of defence (Genton *et al.*, 2005). Therefore, although reductions in both specialists and generalist herbivores have been documented in both introduced ranges, we did not find parallel changes in defence-related traits as predicted by EICA, suggesting such predictions are perhaps too simplistic. Nevertheless, a more detailed survey of herbivory, resistance and the mechanisms of resistance across all three ranges is warranted, particularly given the contrasting patterns of divergence in phenolics identified among the two introduced ranges. Moreover, a more detailed analysis of the alternative functions of these phenolics (e.g., allelopathic interactions and plant structure; Bhattacharya *et al.*, 2010; Li *et al.*, 2010) is required.

A key assumption of EICA is a resource allocation trade-off between defence and growth. However, even when these traits have evolved in the EICA predicted direction, negative genetic correlations have yet to be detected (Franks *et al.*, 2008; Schrieber *et al.*, 2017; Hodgins *et al.*, 2018). Furthermore, a direct trade-off might not be evident as resource reallocation from other traits, drawing from the same resource pool, could allow for the elevated investment in defence related traits and growth simultaneously (Züst & Agrawal, 2017; Hodgins *et al.*, 2018). For instance, an analysis of climate niche shifts in *A. artemisiifolia* has revealed that Eurasian and Australasian ranges on average experience warmer, wetter climates compared to the North American range (van Boheemen *et al.*, 2018). Therefore, reduced investment in abiotic stress tolerance could have allowed for resource reallocation to defence and growth simultaneously. These recently acknowledged complex dynamics underlying competitive ability call for more integrative tests of invasive spread.

### Constitutive versus induced range divergence

We observed a negative association between constitutive and inducible defence-related traits suggesting a trade-off (Koricheva *et al.*, 2004; Agrawal *et al.*, 2010). A decrease in the level and predictability of attack in the introduced range is expected to cause a reduction in constitutive defence and the maintenance or increase in inducible defence (Cipollini *et al.*, 2005; Orians & Ward, 2010; Lande, 2015). In agreement with this prediction constitutive phenolic levels were reduced in Australia, while inducible response did not differ among ranges. Such maintenance of mean inducibility could result from insufficient herbivore pressure, where a selection-drift imbalance could increase inducible variability (Eigenbrode *et al.*, 2008). Although analysis of neutral markers suggests genetic drift has been particularly strong in Australia (van Boheemen *et al.*, 2017), we did not reveal any increase in inducible variation. The growing body of literature testing constitutive versus inducible defence in native and introduced ranges frequently report inconsistent results varying from reductions, to maintenance, to increases in either defence (Cipollini *et al.*, 2005; Eigenbrode *et al.*, 2008; Beaton *et al.*, 2011; Carrillo *et al.*, 2012; Cipollini & Lieurance, 2012; Wang *et al.*, 2012; Wang *et al.*, 2013; Fortuna *et al.*, 2014; Gu *et al.*, 2014; Agrawal *et al.*, 2015; Macel *et al.*, 2017) and calls for more detailed research on the costs-benefit trade-offs of the various responses.

Remarkably, we found evidence of a suppression of phenolics in response to herbivore simulation for some populations, especially those with high constitutive levels, in contrast to some previous studies (Lee *et al.*, 1997; Constabel & Ryan, 1998; Keinänen *et al.*, 2001; Heredia & Cisneros-Zevallos, 2009). Conversely, cardenolide suppression was found in various *Asclepias* species at high constitutive levels (Rasmann *et al.*, 2009), though the mechanistic cause was not discussed (Agrawal *et al.*, 2010). We propose that the retraction of phenolics from damaged leaves could indicate a cost-reducing response when the inducible phenolic compounds have alternative functions (Bhattacharya *et al.*, 2010; Li *et al.*, 2010), or function only in particular aspects of defence response, not induced by the treatment. Similarly, perhaps for those individuals already heavily defended with phenolic compounds increased investment in this chemical defence, which failed to deter an attacking herbivore would have diminishing returns, leading to the potential activation of other defence strategies by the plant. Nevertheless, gaining insight into such cost-benefit associations might prove difficult due to, for instance, issues identifying and addressing all factors influencing the investment of defence-related traits (Neilson *et al.*, 2013).

## Conclusion

We demonstrate divergence of growth and defence traits within multiple introduced ranges that is consistent with rapid adaptation during introduction. Furthermore, we do not find evidence to support the hypothesis that escape from specialist enemies drives the evolution of increased competitive ability in this invasive, as enhanced growth in European populations was not in lieu of defence-related trait reduction. The evolution of growth and defense traits in Australian populations, derived from European founders, occurred rapidly (~80 generations), seemingly unconstrained by strong genetic bottleneck identified in this range (van Boheemen *et al.*, 2017), as measured traits in these two invaded ranges are primarily on opposing ends of the phenotypic spectrum of values. Evidence is growing that adaptation to climate might explain the alarming spread and success of non-natives to a greater extent than release from natural enemies (Colautti *et al.*, 2009; Colautti & Barrett, 2013; Colomer-Ventura *et al.*, 2015). Indeed, we identified repeated latitudinal patterns in phenolics in all three ranges consistent with climate mediated selection, perhaps through corresponding shifts in the biotic community or through direct or indirect selection on phenolics by climate variables. This study emphasizes that intraspecific multi-introduction tests of trait divergence of invasive species provide important insight into contemporary evolutionary process during range expansion.

## Acknowledgements

We would like to thank J. Stephens and A. Wetherhill for sample collection, M. Kourtidou and J. Taylor for greenhouse assistance, K. Nurkowski for genomic analyses and R. Andrew for comments on the MS. A Monash University Dean’s International Postgraduate Research Scholarship was provided to LAB, a Monash University Startup Grant and an ARC grant (DP180102531) to KAH.

## Author contributions

All authors developed the project, with data collection and analyses carried out by LAB and SB, refined by AU and KH. All authors discussed the results, contributed to the MS writing and gave final approval for publication.

## Data accessibility

Sequence data are available at the National Center for Biotechnology Information (NCBI) Sequence Read Archive under Bioproject PRJNA449949.

Scripts are available on https://github.com/lotteanna/defence_adaptation.

Data is available on https://doi.org/10.6084/m9.figshare.8028875.v1

