## Supplement for "Rapid growth and defence evolution following multiple introductions"

### Detailed methods for common garden set-up and data collection

*Common garden set-up*

In 2013-2014, we sampled seeds from 30 maternal plants along transects at each sampling location following the protocol of Hodgins & Rieseberg (2011). We collected plants separated by at least 1-2 meters to reduce the possibility of collecting closely related plants due to limited seed and pollen dispersal. We followed stratification procedures of Willemsen (1975) and stratified at 4°C in sand moistened with 1% plant preservative mixture (ppm) for 6 weeks. Stratification induces germination by imitating winter conditions (Willemsen, 1975). Following stratification, we sowed seeds on damp filter paper in Petri dishes with 1% ppm. We placed dishes in a 30°C germination chamber with 12h light/dark cycle and watered twice daily with 1% ppm to maintain damp filter paper.

After 14-15 days from the start of germination (22^nd^ and 23^rd^ of April, 2015), we randomly selected two seedlings from each mother and planted these into 30-well kwikpot trays containing 100ml Debco Seed Raising Superior Germinating Mix in a random order. We also transplanted two additional seedlings per maternal line, 1-2 days after this first transplant in case focal seedlings died prior to establishment in the greenhouse. Finally, we transplanted late germinating seedlings (sprouting >14 days after we placed seeds in the germination chamber) 5 days after first transplant. We top-watered all plants by hand twice daily and artificially manipulated daylight following the light cycle at the median latitude over all samples (47.3°N) and adjusted timers fortnightly to accommodate the change in day length. Glasshouse temperature was regulated between 20–30 °C. We performed a second transplant of up to three randomly selected seedlings per maternal family (Table S1) on the 22^rd^ to 24^th^ of May, 2015, 1 month after the first transplant, to 100x135mm pots containing 0.7L Debco Seed Raising Superior Germinating Mix and 1.5ml slow-release fertilizer (Osmocote Pro, eight to nine months, 16% N, 4.8% P, 8.3% K, 5.2% S, 1.2% Mg + trace elements). To explore constitutive defence, we selected one seedling from four maternal lines originating from 28 North American, 32 European and 20 Australian locations (Table S1). A separate greenhouse experiment was conducted to test whether the inducibility of defence response varied among plant origins (hereafter, “induction experiment”). We used a subset of populations used in the constitutive experiment (10 North American, 17 European and 12 Australian locations, Table S1). For each population, we selected four maternal lines, and grew two seedlings per line as above. One seedling per mom was allocated to either the control or simulated herbivory treatment. We simulated herbivory by vertically cutting off half of the newest fully formed leaf (wounding) and subsequently spraying the whole plant with 1mM methyl jasmonate (MeJA) (Campos‐Vargas & Saltveit, 2002; Heredia & Cisneros-Zevallos, 2009). Control plants were not wounded and were sprayed with distilled water. We watered plants twice daily and randomized tray locations weekly.

*Genetic data collection and analyses*

Four weeks after second transplant, we harvested a 5-7 cm young leaf for DNA extraction and sealed these inside 1.5mL Eppendorf tubes, which were flash-frozen in liquid nitrogen. We stored samples on dry ice before placing them in an -80 °C freezer. We extracted DNA from 29-99mg (mean=72.5mg) of leaf tissue of 861 individuals (84 populations) from the control treatment using the Glass Fiber Plate DNA Extraction Protocol (CCDB, Guelph, ON, Canada) and assessed DNA quantity (>8.5 ng/μl) using a QuBit broad-sensitivity DNA quantification system (Invitrogen, Carlsbad, CA, USA). We performed double-digest genotype-by-sequencing library preparation We added 200 ng of high-quality DNA in 7.2 μL water to 2.0 uL CutSmart Buffer 10x, 0.4 uL Pst-1 HF (NEB), 0.4 uL Msp1. We digested samples for 8h at 37°C, 20 minutes at 65°C. To each reaction, we added 2.0 uL 10x CutSmart Buffer, 4.0 uL 10mM ATP, 0.5uL T4 DNA Ligase, 8 uL H2O, 1uL 10mM common adaptor and 5uL 0.6ng/uL barcoded adaptor. We ligated samples for 3h at 22°C and 20 minutes at 65°C. We mixed all samples with 6144 uL Sera-Mag beads (Thermo Fisher). After 15-minute incubation at room temperature, we allotted samples to 7 1.5mL tubes and placed these in Dyna-Mag 2 (Thermo Fisher) magnet for 4 minutes. We removed clear liquid and washed 3 times using 80% EtOH and once with 100% EtOH. We eluted in 150 uL 10 mM Tris pH 8.0. We amplified 8 reactions each with 3uL of elution and 7.5uL H2O, 12.5 μL KAPA 2x MasterMix, 1uL of 12.5mM each PCR primers f&r. We cycled reactions at 98°C for 1 minute, followed by 20s at 62°C and 30s at 72°C. Following 16 cycles we additionally kept samples at 72°C for 5 minutes. After amplification, we cleaned up 30 μL from each well using the Bioline PCR and Gel kit (Bioline). We eluted the purified product in 30uL buffer. We performed a size selection by running the cleaned PCR product on a 2% agarose gel and removing the 400-600bp fragment. This gel fragment was cleaned up using the Bioline PCR and Gel Kit (Biolin1) and eluted in 20 uL H2O. The eleven ddGBS libraries were paired-end sequenced by the Biodiversity Sequencing Centre at UBC on the Illumina HiSeq 2000 platform, two libraries per lane.

We aligned and filtered raw sequences following van Boheemen *et al.* (2017a). Briefly, SNPs were aligned using BWA-mem (Li & Durbin, 2009) to a draft reference genome for *A. artemisiifolia* (van Boheemen *et al.*, 2017a). We called variants with GATK UnifiedGenotyper (McKenna *et al.*, 2010) and filtered SNPs as follows: quality threshold of a Q-score ≥50; a minimum quality by depth of 2, a maximum Fisher-Strand bias of 60.0, minimum mapping quality rank sum test of -12.5, minimum root mean square mapping quality of 40.0, and a minimum read position rank sum test of -8.0; genotype and variant quality of ≥20, depth of 5-240 and a minor allele frequency of 0.05. We identified a total of 11,598 polymorphic biallelic SNPs with 50% SNP call rate. We calculated individual heterozygosity (H_O_) as the proportion of heterozygous loci out of the total number of called genotypes for each individual across 836 control treatment plants and excluded individuals with >80% non-called genotypes (25 out of 861 individuals).

We inferred population genetic structure with STRUCTURE v2.3.4, a Bayesian clustering method that allocates individuals into clusters on the basis of their genotypes (Pritchard *et al.*, 2000). We used this program to calculate population q-scores for the most likely K, used in subsequent analyses to correct for population structure. From the 11,598 SNPs identified, we selected 1024 unlinked SNPs by shuffling the full SNP table and randomly drawing one SNP from each contig. We ran STRUCTURE on these SNPs for each range, using the admixture model, correlated allele frequencies, no location prior for the number of clusters (K) ranging from 1 to 10, with 20 independent runs per K. We sampled from a uniform prior for alpha, whilst allowing for alpha to vary between clusters, accounting for unequal sample sizes (Wang, 2017) Each run comprised of a burn-in of 200,000 followed by 1,000,000 iterations. We used log probability and delta K statistic to determine the uppermost clustering level (Evanno *et al.*, 2005). We used CLUMPAK (Kopelman *et al.*, 2015) to test for multimodality (non present) for the most likely K (=2).

**Table S1.** Sampling locations within the native North American and introduced European and Australian *Ambrosia artemisiifolia* ranges, including state or province (North America and Australian) and country (North America & Europe), population ID, geographic coordinates (WGS84) and if sampling location was included in the constitutive (one seedling per maternal line) and/or inducible (two seedlings per maternal line) experiment.

| **Range** | **Population ID** | **State/Country** | **Latitude** | **Longitude** | **Constitutive** | **Inducible** |
| --- | --- | --- | --- | --- | --- | --- |
| North America, native | AL | AL, USA | 30.6754 | -87.5908 | yes |  |
|  | MP | MS, USA | 31.2078 | -89.0664 | yes | yes |
|  | GA | GA, USA | 31.6343 | -81.4069 | yes | yes |
|  | AR | AR, USA | 33.9755 | -91.4134 | yes | yes |
|  | SC | SC, USA | 34.2252 | -81.3431 | yes | yes |
|  | NC | NC, USA | 35.6070 | -83.0217 | yes |  |
|  | TN | TN, USA | 36.2678 | -86.4606 | yes | yes |
|  | MO | MO, USA | 37.0064 | -94.3501 | yes | yes |
|  | KY | KY, USA | 38.6259 | -85.0074 | yes |  |
|  | KS | KS, USA | 38.6857 | -96.4927 | yes | yes |
|  | NB | NE, USA | 40.0439 | -96.3314 | yes | yes |
|  | OH | OH, USA | 40.4876 | -82.7270 | yes | yes |
|  | PA | PA, USA | 40.9659 | -78.1748 | yes |  |
|  | NY | NY, USA | 41.4415 | -74.5291 | yes |  |
|  | MS | MA, USA | 42.0883 | -72.0962 | yes |  |
|  | IW | IW, USA | 42.6780 | -96.5024 | yes | yes |
|  | AA8 | MN, USA | 44.3254 | -95.9583 | yes | yes |
|  | MN2 | ON, Canada | 44.4472 | -79.8039 | yes | yes |
|  | MA | ME, USA | 44.7712 | -68.9709 | yes |  |
|  | WI | WI, USA | 44.8793 | -89.4238 | yes | yes |
|  | ON3 | ON, Canada | 45.3297 | -74.8933 | yes | yes |
|  | ON2 | ON, Canada | 45.7445 | -77.0294 | yes |  |
|  | NB1 | NB, Canada | 45.8786 | -66.9785 | yes | yes |
|  | AA5 | MN, USA | 46.2171 | -96.0502 | yes | yes |
|  | MI | MI, USA | 46.3581 | -84.8807 | yes | yes |
|  | QC2 | QC, Canada | 46.8867 | -70.8684 | yes |  |
|  | QC3 | QC, Canada | 47.6788 | -69.0220 | yes |  |
|  | AA2B | MN, Canada | 49.8378 | -97.3293 | yes | yes |
| Europe, introduced | EU35 | Serbia | 43.0918 | 21.9380 | yes |  |
|  | EU36 | Bulgaria | 43.3152 | 24.2598 | yes | yes |
|  | EU34 | Bulgaria | 43.4704 | 25.6707 | yes | yes |
|  | EU37 | Serbia | 43.9179 | 20.7331 | yes | yes |
|  | EU10 | France | 43.9324 | 4.3205 | yes |  |
|  | EU32 | Romania | 44.1627 | 28.5098 | yes |  |
|  | EU33 | Romania | 44.4046 | 26.1348 | yes | yes |
|  | EU09 | Italy | 45.0654 | 7.5923 | yes |  |
|  | EU11 | France | 45.0744 | 4.7509 | yes | yes |
|  | EU31 | Romania | 45.3740 | 27.0720 | yes | yes |
|  | EU07 | Italy | 45.47089 | 8.93683 |  | yes |
|  | EU06 | Italy | 45.5707 | 8.7855 | yes | yes |
|  | EU27 | Romania | 45.6870 | 25.6581 | yes | yes |
|  | EU38 | Croatia | 45.7153 | 15.6538 | yes |  |
|  | EU08 | Switzerland | 45.9309 | 8.9838 | yes | yes |
|  | EU01 | Slovenia | 46.0361 | 15.2961 | yes |  |
|  | EU15 | France | 46.0522 | 5.3351 | yes |  |
|  | EU16 | Switzerland | 46.1622 | 6.0094 | yes | yes |
|  | EU26 | Romania | 46.2370 | 24.8540 | yes |  |
|  | EU12 | France | 46.6643 | 4.3278 | yes |  |
|  | EU13 | France | 46.8003 | 4.9724 | yes | yes |
|  | EU02 | Hungary | 47.1309 | 16.9031 | yes |  |
|  | EU03 | Hungary | 47.3278 | 19.7307 | yes | yes |
|  | EU14 | France | 47.4556 | 5.2121 | yes |  |
|  | EU04 | Slovakia | 47.8799 | 18.1545 | yes | yes |
|  | EU25 | Romania | 47.9773 | 23.0443 | yes |  |
|  | EU24 | Slovakia | 48.4892 | 21.8062 | yes |  |
|  | EU22 | Czech | 49.4180 | 17.9615 | yes | yes |
|  | EU30 | Poland | 49.8685 | 23.0118 | yes |  |
|  | EU21 | Czech | 50.1900 | 15.0604 | yes | yes |
|  | EU23 | Poland | 50.4430 | 18.8634 | yes |  |
|  | EU17 | the Netherlands | 51.1200 | 5.8403 | yes | yes |
|  | EU20 | Germany | 51.6328 | 14.1844 | yes |  |
| Australia, introduced | AU01 | NSW | -35.6411 | 150.1274 | yes |  |
|  | AU32 | NSW | -31.4721 | 152.6836 | yes |  |
|  | AU33 | NSW | -31.4419 | 152.4651 | yes | yes |
|  | AU29 | NSW | -30.9195 | 152.5825 | yes |  |
|  | AU30 | NSW | -30.7406 | 152.9150 | yes | yes |
|  | AU03 | NSW | -30.3881 | 152.9582 | yes | yes |
|  | AU27 | NSW | -30.2413 | 152.5855 | yes |  |
|  | AU26 | NSW | -30.0514 | 152.9849 | yes | yes |
|  | AU04 | NSW | -29.6310 | 153.0368 | yes |  |
|  | AU24 | NSW | -29.4358 | 152.3848 | yes | yes |
|  | AU23 | NSW | -28.9263 | 152.3740 | yes |  |
|  | AU09 | NSW | -28.8688 | 151.1670 | yes |  |
|  | AU05 | NSW | -28.7668 | 153.3966 | yes |  |
|  | AU21 | NSW | -28.3870 | 152.6099 | yes | yes |
|  | AU19 | QLD | -28.0122 | 153.1677 | yes |  |
|  | AU18 | QLD | -27.7850 | 153.2751 | yes |  |
|  | AU15 | QLD | -27.3792 | 152.8016 | yes | yes |
|  | AU12 | QLD | -26.8858 | 152.1369 | yes |  |
|  | AU13 | QLD | -26.3913 | 152.7937 | yes |  |
|  | AU11 | QLD | -25.3655 | 152.9156 | yes | yes |

**Table S2.** *Ambrosia artemisiifolia* defence-related trait responses model adjusted means (standard error) for each range (corresponds to Table 1 & Fig. 1, main text).

|  | North America | Europe | Australia |
| --- | --- | --- | --- |
| Phenolic concentration | 14.524(0) | 15.824(0) | 10.928(0) |
| Phenolic richness | 40.72(0.869) | 43.503(1.09) | 32.699(1.935) |
| Trichome density | 99.893(4.607) | 97.973(4.226) | 113.437(6.524) |

**Table S3.** *Ambrosia artemisiifolia* model adjusted means (standard error) for each range in the inducible experiment in multivariate (individual phenolic compounds) and univariate analyses (corresponds to Table 2 & Fig. 3, main text).

| Range | Native | | Europe | | Australia | |
| --- | --- | --- | --- | --- | --- | --- |
| Treatment | Control | Wounding + MeJa | Control | Wounding + MeJa | Control | Wounding + MeJa |
| Phenolic concentration | 52.803 (10.406) | 44.618 (10.407) | 69.724 (10.404) | 61.539 (10.405) | 19.949 (10.402) | 11.764 (10.403) |
| Phenolic richness | 24.06 (1.934) | 23.261 (1.943) | 30.616 (2.18) | 29.816 (2.189) | 18.569 (3.359) | 17.77 (3.354) |

**Table S4.** Constitutive defence trait response of *Ambrosia artemisiifolia* individuals, model adjusted means (standard error) for each range (corresponds to Table 3 & Fig. 4, main text).

|  |  | Figure 4 | North America | Europe | Australia |
| --- | --- | --- | --- | --- | --- |
| Shoot biomass | Phenolic concentration | A | 13.881(0) | 16.502(0.001) | 8.246(0) |
|  | Phenolic richness | B | 40.985(0.764) | 42.671(0.751) | 33.863(1.133) |
|  | Trichome density | C | 97.429(20.172) | 89.073(18.13) | 126.099(39.006) |
| Specific leaf area | Phenolic concentration | D | 13.581(0.001) | 15.313(0) | 10.367(0.001) |
|  | Phenolic richness | E | 40.345(0.967) | 41.182(0.817) | 36.106(1.325) |
|  | Trichome density | F | 99.412(26.303) | 96.815(21.298) | 113.845(42.536) |
